# Enhanced intestinal reabsorption due to reduced metabolism of ceftriaxone

**DOI:** 10.1101/205880

**Authors:** Tapas K Sar, Indranil Samanta, Rabindra N Hansda, Rinku Buragohain, Tapan K Mandal

## Abstract

The polyherbal drug (Fibrosin^®^) is often used as supportive therapy with intramuscular ceftriaxone injection for treatment of mastitis. A single dose of ceftriaxone at 50 mg kg^−1^ was administered intramuscularly in six healthy lactating goats and a single oral dose of Fibrosin^®^ (1.9 gm) was given 1 hr prior to intramuscular ceftriaxone injection to study disposition of ceftriaxone. Plasma and urine samples were collected at predetermined time schedule and ceftriaxone/ceftizoxime was extracted and analyzed by HPLC. Ceftriaxone persisted for 3 hr in plasma of fibrosin treated healthy lactating goats. Mean t_½_ K (distribution half life) following absorption phase and t_½_ K′(elimination half life) following intestinal re-absorption phase were respectively, 0.091 ± 0.01 and 0.43 ± 0.03 hr with a re-absorption half life (t_½_Ka′) of 0.18 ± 0.003 hr.Ceftriaxone at a lower concentration (67.91 ± 9.42 μg ml^−1^) was recovered at 24 hr post dosing from urine. This is the first report of pharmacokinetic interaction of intramuscular ceftriaxone injection and the oral polyherbal drug (which was found to be a cytochrome P_450_ inhibitor).

## Introduction

Ceftriaxone, a third generation cephalosporin, is active against a wide range of gram negative and gram positive organisms. The drug is converted in liver to a major active metabolite i.e. ceftizoxime in goat (Sar *et al*, 2006; Sar *et al*, 2008; Sar *et al*, 2013) and cow (Sar *et al*, 2010). Ceftizoxime, a broad spectrum third generation cephalosporin, is effective against a wide variety of aerobic/anaerobic gram positive and gram negative bacteria and remains highly stable in presence of β lactamases. The previous studies by our team suggested a typical absorption-reabsorption pattern of ceftriaxone in plasma of goat following intramammary administration at 50 mg kg^−1^ with 1 hr prior oral dosing of the polyherbal drug (Fibrosin^®^) (Sar *et al*, 2011) and after intramuscular administration of ceftriaxone only at 50 mg kg^−1^ (Sar *et al*, 2013). Reabsorption of ceftriaxone was also observed following intravenous injection of ceftriaxone at 50 mg kg^−1^ in lactating goats with 1 hr prior oral dosing of the polyherbal drug (Fibrosin^®^) for which ceftriaxone was again available from 36 to 84 hr post dosing in plasma following its disappearance from 3 hr post dosing (Sar *et al* 2006). However, reabsorption of ceftriaxone could not be observed after intravenous injection of ceftriaxone only at 50 mg kg^−1^ (Sar *et al*, 2006; Guerrini *et al*, 1983). Administration of ceftriaxone at 46 mg kg^−1^ intravenously to sheep resulted in more rapid non-renal elimination of ceftriaxone and biliary secretion of ceftriaxone into the duodenum was reported (Arvidsson *et al*, 1982; Stoeckel *et al*, 1981). But, reabsorption may also occur in intestine. Although the previous studies on intramuscular pharmacokinetics of ceftriaxone in cow calves (Soback *et al*, 1988; Johal &Srivastava, 1998), camel (Goudah, 2008), cat (Abarellos *et al*, 2007) and goat ( Ismail *et al*, 2005) did not reveal any absorption-reabsorption pattern but probably blood collection at frequent interval of time may show the two phases. The veterinarians frequently use ceftriaxone intramuscularly for treatment of susceptible bacterial infections. Fibrosin^®^, a polyherbal drug marketed by Legend Remedies Pvt. Ltd.,Vadadora, India, is used as supportive therapy for mastitis. The polyherbal drug facilitates cleaning of udder by clearing the tissue debris and helps in down flow of milk in mastitis (leaflet of Fibrosin^®^). Fibrosin^®^ inhibited cytochrome P_450_ in liver of goats following single oral dosing at 1.9 gm, but intravenous ceftriaxone injection at 50 mg kg^−1^ with 1 hr prior oral dosing of Fibrosin^®^ caused induction of cytochrome P_450_ in liver of goats (Sar *et al*, 2006). Fibrosin^®^ enhanced the penetration of ceftizoxime through milk-blood barrier and increased the bioavailability of ceftizoxime in milk following a single dose intramammary administration of ceftriaxone at 50 mg kg^−1^ after its metabolism in liver (Sar *et al*, 2011). Oral administration of Fibrosin^®^ also could be preferred with concurrent administration of intramuscular ceftriaxone for treatment of sensitive bacterial infections except mastitis as it causes zero level milk residue of ceftizoxime (Sar *et al*, 2014). The polyherbal drug also returned increased milk alkaline phosphatase and catalase activity in mastitic goats to normal level and maintained normal glutathione level with significantly increased lactoperoxidase activity with 1 hr prior oral administration to single intravenous dosing of ceftriaxone (Sar *et al*, 2012). The polyherbal drug also protected mammary gland from tissue damage caused by intramammary administration of ceftriaxone (Sar *et al*, 2015). Milk alkaline phosphatase and catalase activity and reduced glutathione level did not differ significantly between healthy and fibrosin treated goats following single intramuscular dosing of ceftriaxone without and with 1 hr prior oral administration of Fibrosin^®^ (Sar *et al*, 2014). Disposition of ceftriaxone exhibited absorption-reabsorption pattern in healthy goats, while it was absent in carbontetrachloride treated hepatopathic and no reabsorption in uranyl nitrate treated nephropathic goats following single intramuscular and intravenous dosing and at50 mg kg^−1^ (Sar *et al*, 2013). Ceftriaxone was reported to induce cytochrome P_450_ in liver and the polyherbal drug (Fibrosin^®^) was found to inhibit cytochrome P_450_ (Sar *et al*, 2006). Disposition of ceftriaxone in healthy goats following single intramuscular dosing at 50 mg kg^−1^ had already been reported (Sar *et al*, 2013).

Therefore, the present study was aimed to evaluate disposition kinetics of ceftriaxone in presence of the polyherbal drug (Fibrosin^®^) and to observe its effect on intestinal reabsorption of ceftriaxone following single intramuscular administration of ceftriaxone with 1 hr prior oral dosing of Fibrosin^®^ as the polyherbal drug was found to inhibit cytochrome P_450_ oxidase system in liver. The study was also conducted to explore possible pharmacokinetic interactions between intramuscular ceftriaxone injection and any cytochrome P_450_ inhibitor drug.

## Results

The recovery percentage of ceftriaxone and ceftizoxime were 81.14±4.95% and 80.37±2.30%, respectively in plasma. In urine, the recovery percentage of ceftriaxone and ceftizoxime were 85.83±4.21% and 87.56±2.84%, respectively. The chromatograms of standard ceftriaxone and ceftizoxime were presented in Figure 1(a) and Figure 1(b), blank plasma in Figure 2, chromatograms of ceftriaxone and ceftizoxime extracted from plasma in Figure 3(a) and Figure 3(b), standard curve of ceftriaxone and ceftizoxime in plasma in Figure 4.

**Fig. 1(a).**
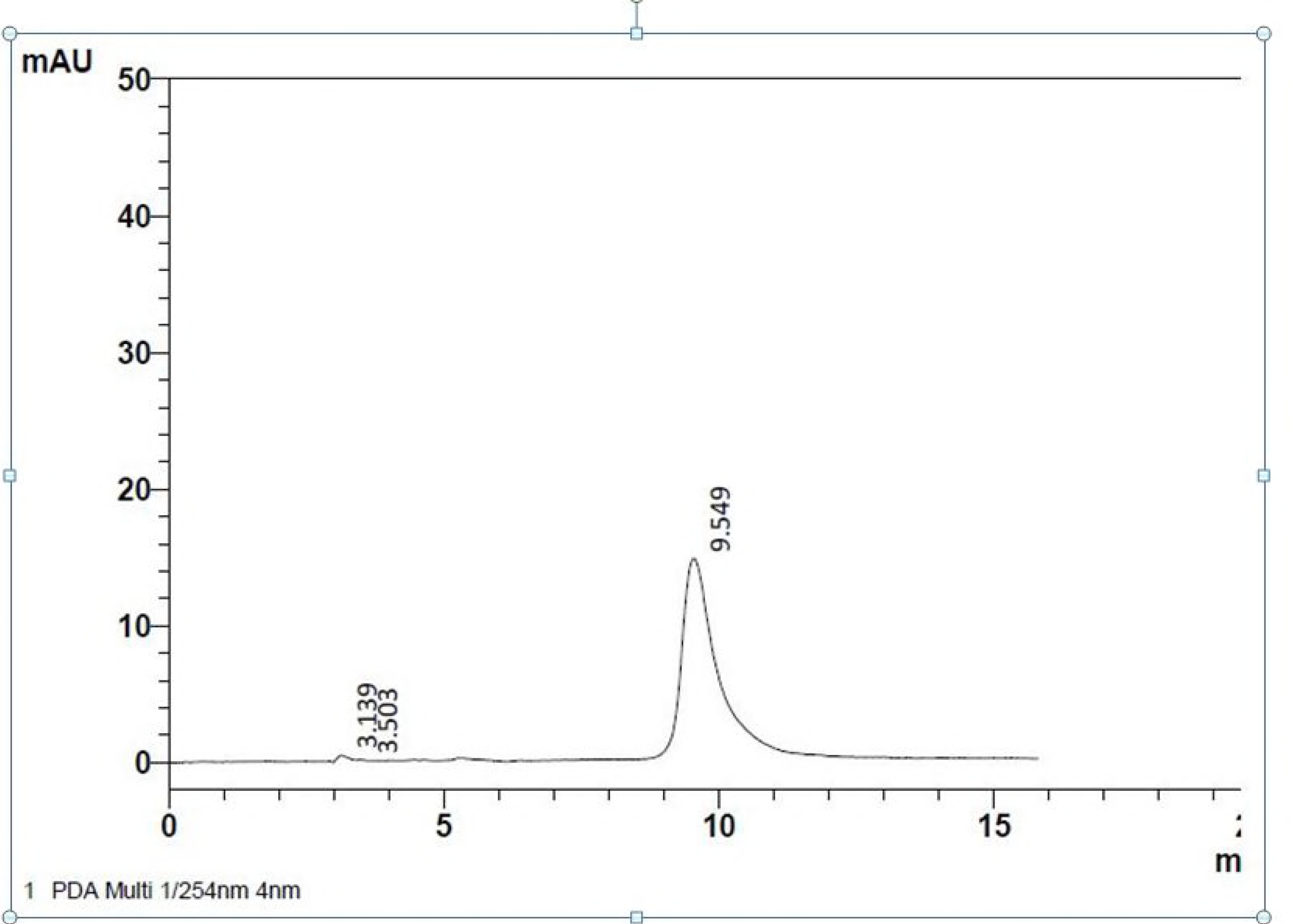
Chromatogram of ceftriaxone (10 ppm); RT=9.549

**Fig. 1(b).**
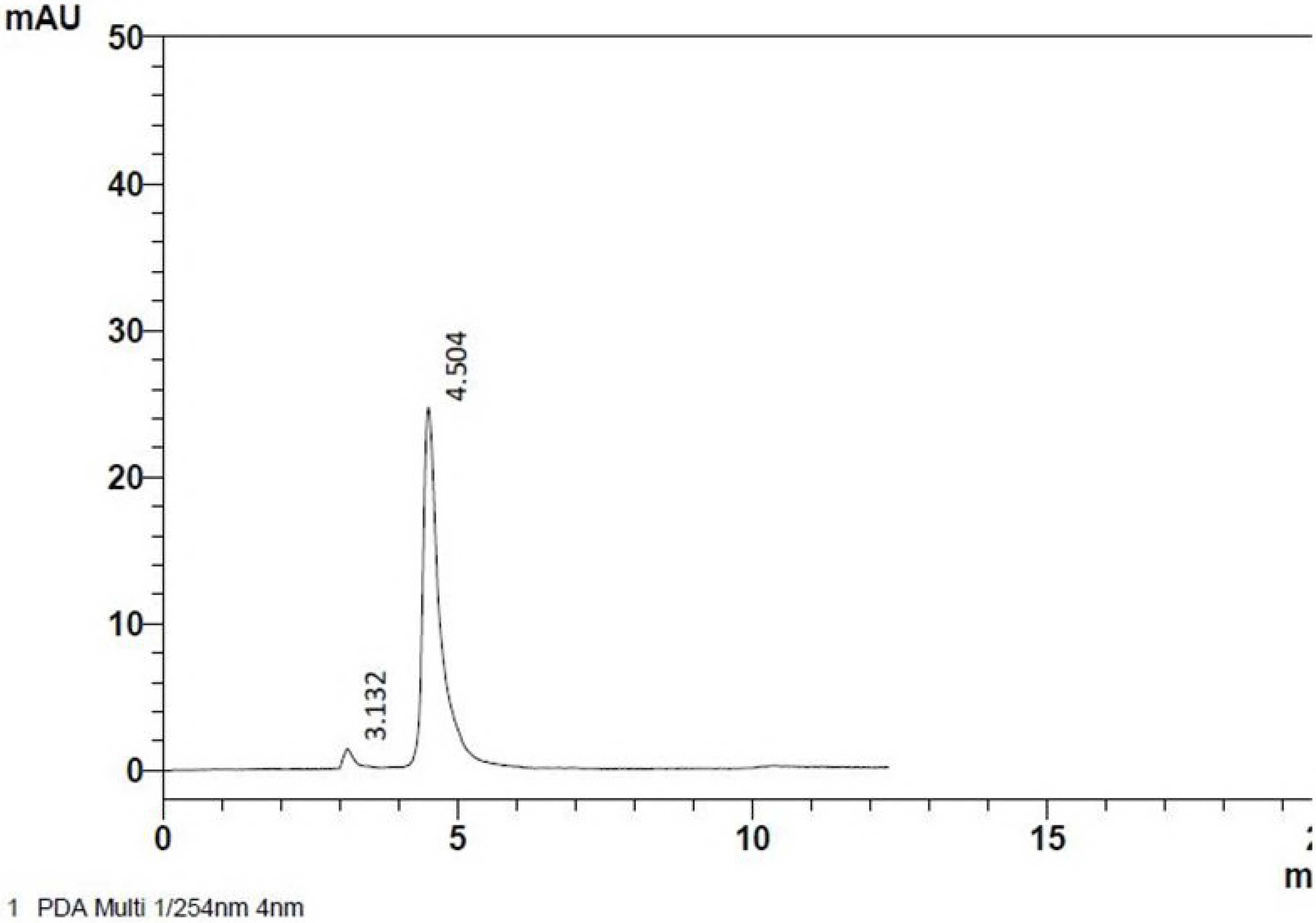
Ceftizoxime (10 ppm); RT = 4.504

**Fig. 2.**
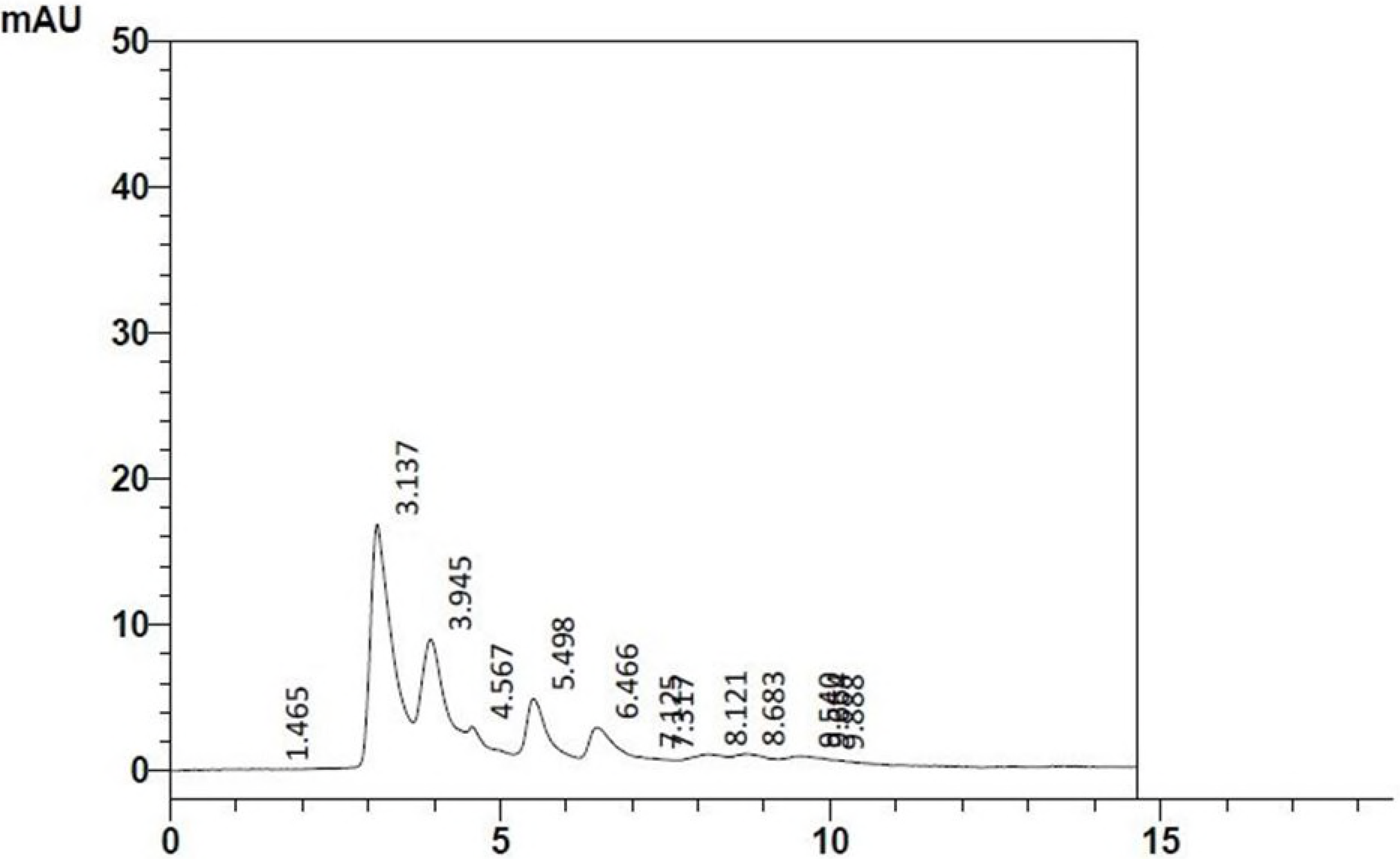
Chromatogram of blank plasma

**Fig. 3(a).**
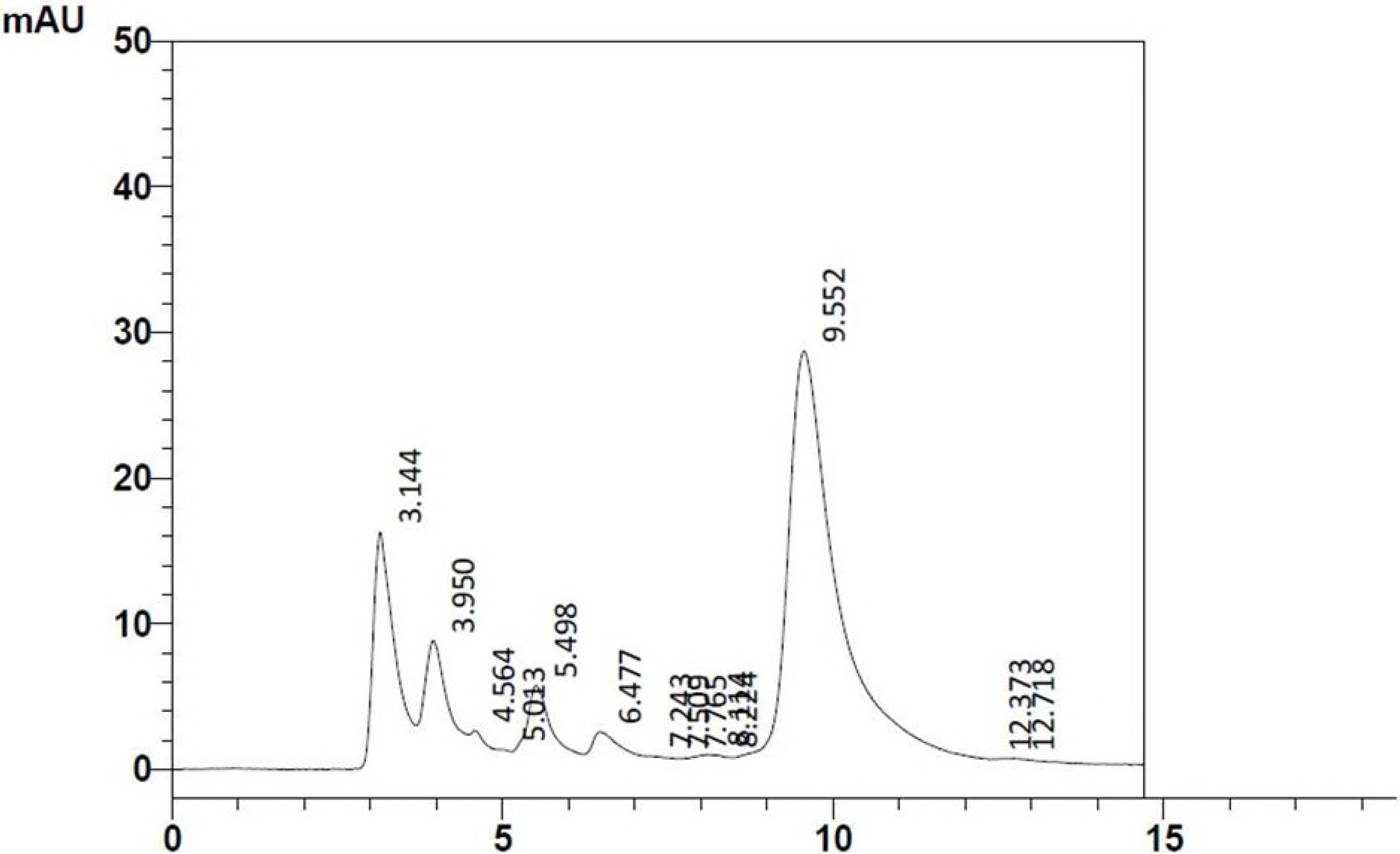
Chromatogram of ceftriaxone extracted from plasma

**Fig. 3(b).**
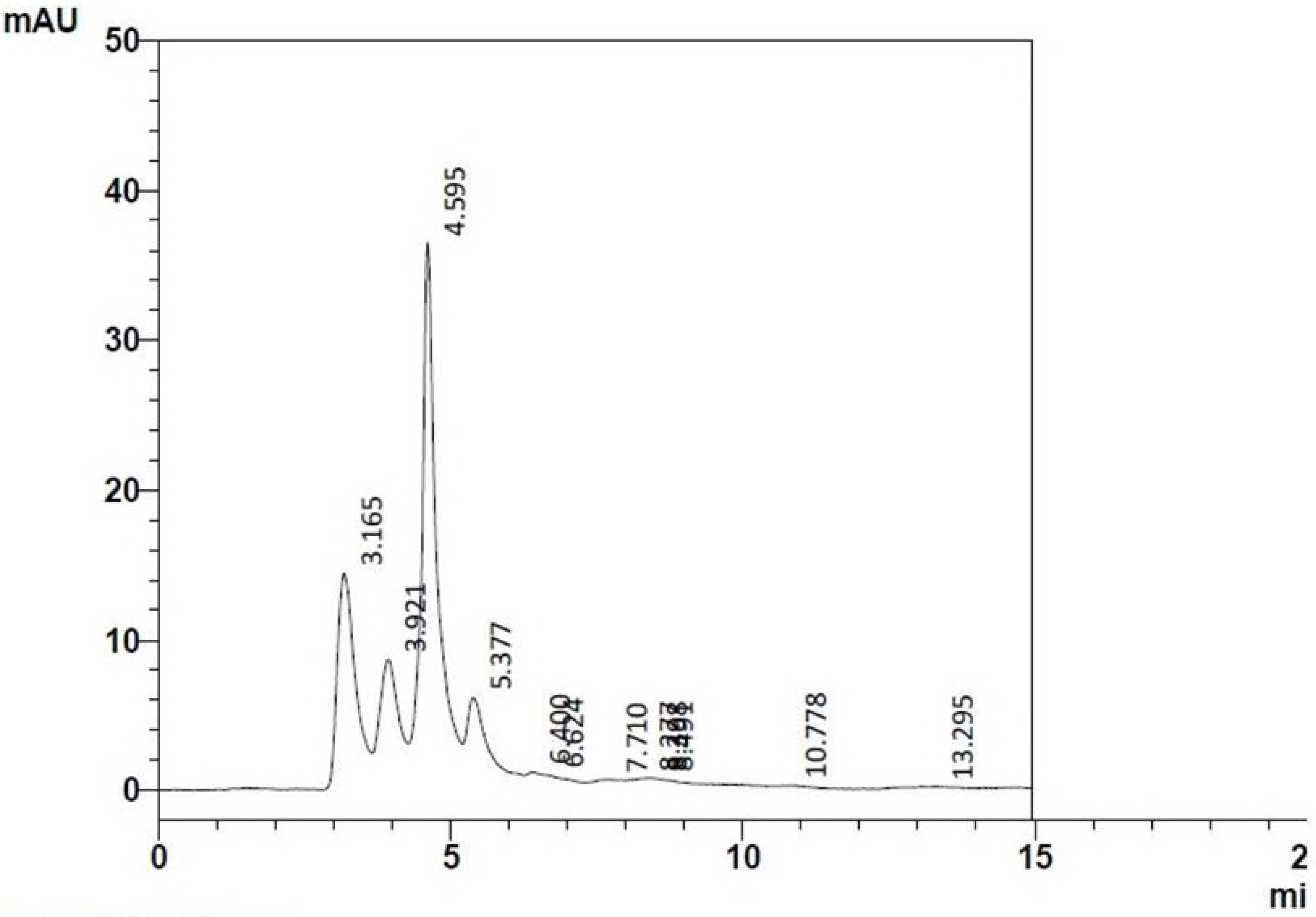
Chromatogram of ceftizoxime extracted from plasma (*in-vitro*)

**Fig.4.**
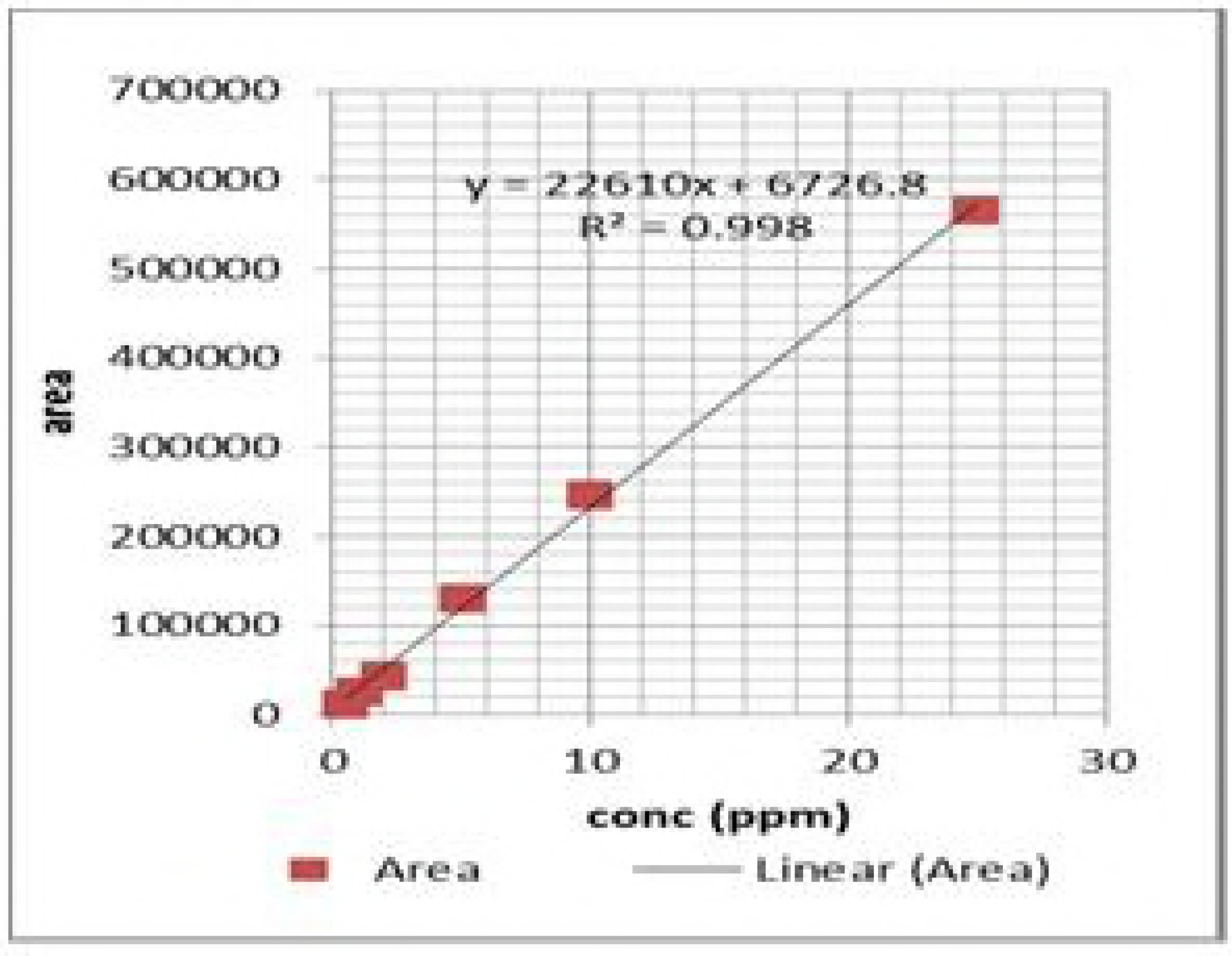
Standard curve of recovery of ceftriaxone (a) and ceftizoxime (b) in plasma

The effects of Fibrosin^®^ on liver function tests were presented in Table 1.The normal icterus index, serum bromosulfophthalein clearance (BSP), serum aspartate transaminase activity (AST) and serum alanine transaminase activity (ALT) were 4.64^NS^ ± 0.10 unit, 3.32 ^NS^ ± 0.36 min, 340.00 ^NS^ ± 11.64 μg pyruvic acid/mL/h and 92.00 ^NS^ ± 7.83 μg pyruvic acid/mL/h, respectively in healthy lactating goats which did not differ significantly on day 3 and 7 after single dose oral administration of Fibrosin^®^ (1.9 gm).

**Table 1.** Mean values with SE of icterus index, serum bromosulfophthalein clearance (BSP), serum aspartate transaminase activity (AST), serum alanine transaminase activity (ALT) in healthy lactating goats following single dose oral administration of fibrosin^®^ (1.9gm).

| Day | Icterus index (unit) | BSP Clearance (t<sub>1/2</sub> min) | AST activity (µg pyruvic acid/mL/h) | ALT activity (µg pyruvic acid/mL/h) |
| --- | --- | --- | --- | --- |
| 0 | 4.64<sup>NS</sup> ± 0.10 | 3.32<sup>NS</sup> ± 0.36 | 340.00<sup>NS</sup> ± 11.64 | 92.00<sup>NS</sup> ± 7.83 |
| 3 | 4.61<sup>NS</sup> ± 0.09 | 3.05<sup>NS</sup> ± 0.15 | 340.00<sup>NS</sup> ± 11.67 | 85.33<sup>NS</sup> ± 7.73 |
| 7 | 4.30<sup>NS</sup> ± 0.07 | 3.07<sup>NS</sup> ± 0.11 | 331.67<sup>NS</sup> ± 7.88 | 83.33<sup>NS</sup> ± 6.68 |

Means present in each column do not differ significantly

Mean plasma concentration with SE of ceftriaxone in healthy lactating goats with 1 hr pre single dose oral administration of fibrosin (1.9 gm) after single dose intramuscular administration at 50 mg kg^−1^ have been incorporated in (Table 2). Ceftriaxone attained a plasma concentration of 38.50 ± 5.34 μg ml^−1^ at 0.08 hr which started to increase gradually with a peak concentration of 69.00 ± 8.08 μg ml^−1^ at 0.33 hr and declined at 0.50 hr (19.50 ± 5.34 μg ml^−1^) which again started to increase from 0.66 hr post dosing, achieved a maximum concentration of 72.50 ± 7.21 μg ml^−1^ at 1 hr and declined to a minimum concentration of 3.00 ± 1.01 μg ml^−1^ at 3 hr post dosing in lactating goats treated with fibrosin.

**Table 2.** Mean plasma concentration (μg ml^−1^) of ceftriaxone in healthy lactating goats without and with 1 hr pre single dose oral administration of fibrosin (1.9 gm) after single dose intramuscular administration at 50 mg kg^−1^

| Time (hr) | Fibrosin treated healthy lactating goats |
| --- | --- |
| 0.08 | $38.50^{\text{bcd}} \pm 5.34$ |
| 0.16 | $46.90^{\text{cdef}} \pm 6.40$ |
| 0.25 | $56.00^{\text{def}} \pm 6.92$ |
| 0.33 | $69.00^{\text{ef}} \pm 8.08$ |
| 0.50 | $19.50^{\text{abc}} \pm 5.34$ |
| 0.66 | $29.80^{\text{abcd}} \pm 6.37$ |
| 0.83 | $45.66^{\text{bcde}} \pm 7.41$ |
| 1 | $72.50^{\text{f}} \pm 7.21$ |
| 2 | $14.90^{\text{ab}} \pm 3.69$ |
| 3 | $3.00^{\text{a}} \pm 1.01$ |
| 4 | BDL |

BDL – Below detection limit; Means bearing a common superscript do not differ significantly ( $p < 0.05$ )

The semilogarithmic plot of mean plasma level time profile and disposition kinetic parameters of ceftriaxone in healthy lactating goats after single dose intramuscular administration at 50 mg kg^−1^ with 1 hr pre single dose oral administration of Fibrosin^®^ have been presented in Fig.5 and Table 3. In healthy fibrosin treated lactating goats, the values of K and t_½_K of distribution phase were 7.85 ± 1.07 hr^−1^ and 0.091 ± 0.01 hr respectively, while K′ and t½ K′ values of elimination phase were respectively 1.65 ± 0.15 hr^−1^ and 0.43 ± 0.035 hr. The values of Ka and t_½_Ka of absorption phase were respectively 10.75 ± 0.80 hr^−1^ and 0.064 ± 0.004 hr. But, the values of Ka′ and t_½_Ka′ of reabsorption phase were found to be 3.86 ± 0.10 hr^−1^ and 0.18 ± 0.003 hr. Mean t_½_Ka′ of reabsorption phase was manifold higher compared to mean t_½_Ka of absorption phase indicating prolonged and enhanced reabsorption of ceftriaxone compared to its absorption. Mean AUC and Vdarea values were 247.50 ± 24.89 μg hr ml^−1^ and 0.13 ± 0.003 L kg^−1^ that indicated wider distribution coverage with lesser volume of distribution.

**Fig.5.**
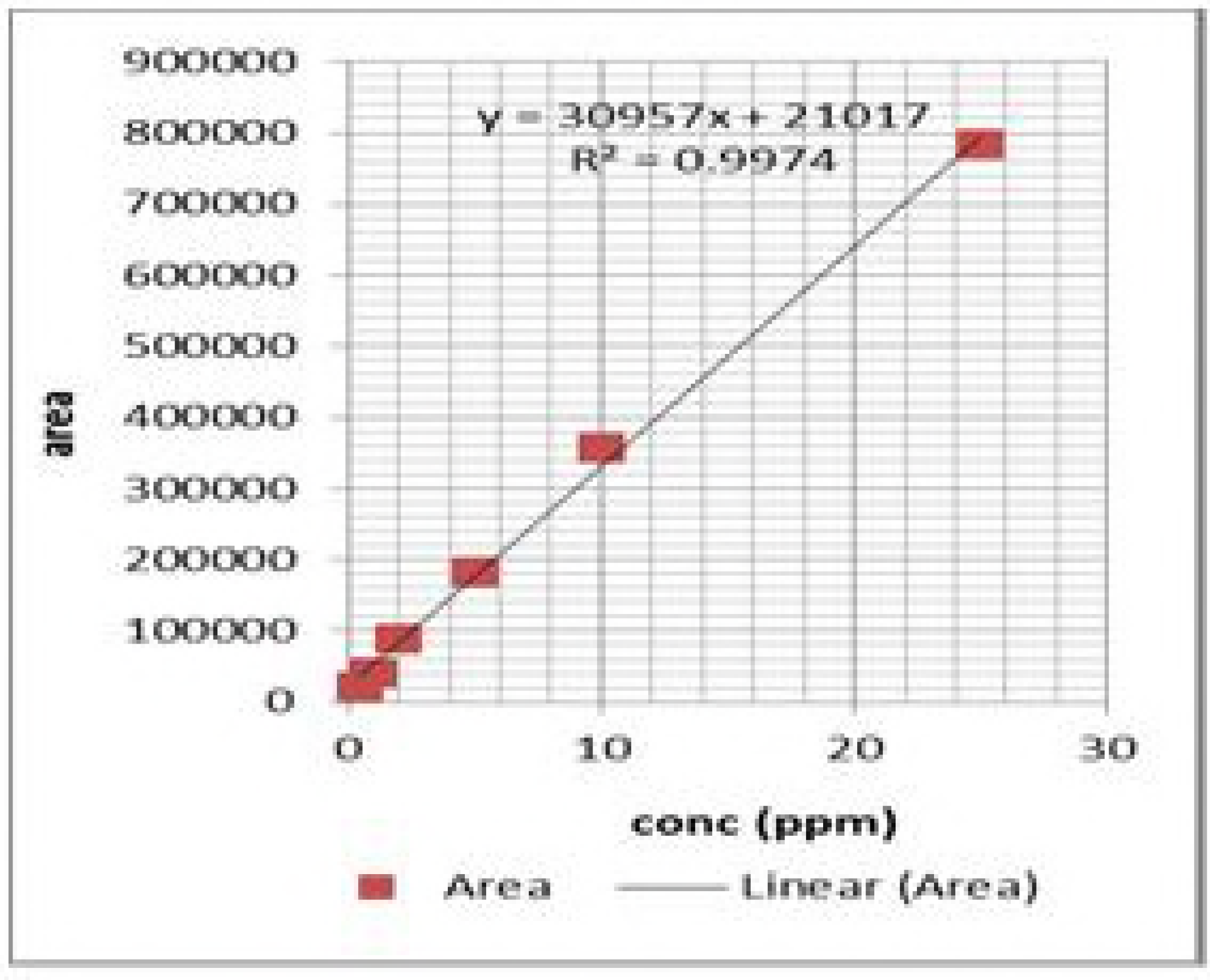
Semilogarithmic plot of mean plasma concentration (μg/ml) of ceftriaxone in lactating goats after single-dose intramuscularadministration of ceftriaxone at 50mg/kg with 1 hr pre single dose oral administration of Fibrosin^®^ [X-axis: time (hr), Y-axis: plasma concentration of ceftriaxone (μgm ml^−1^)].

**Table 3.** Mean kinetic parameter of ceftriaxone in healthy lactating goats without and with 1 hr pre single dose oral administration of fibrosin (1.9 gm) after single dose intramuscular administration at 50 mg kg^−1^

| Kinetic parameters | Fibrosin treated healthy lactating goats |
| --- | --- |
| K<sub>a</sub> (hr<sup>-1</sup>) | 10.75 ± 0.80 |
| t<sub>1/2</sub>k<sub>a</sub> (hr) | 0.064 ± 0.004 |
| K (hr<sup>-1</sup>) | 7.85 ± 1.07 |
| t<sub>1/2</sub>k (hr) | 0.091 ± 0.01 |
| K'<sub>a</sub> (hr<sup>-1</sup>) | 3.86 ± 0.10 |
| t<sub>1/2</sub> K'<sub>a</sub> (hr) | 0.18 ± 0.003 |
| K' (hr<sup>-1</sup>) | 1.65 ± 0.15 |
| t<sub>1/2</sub>k' (hr) | 0.43 ± 0.035 |
| Vd<sub>area</sub> (L kg<sup>-1</sup>) | 0.13 ± 0.003 |
| AUC (µg hr ml<sup>-1</sup>) [Based on trapizoid method] | 247.50 ± 24.89 |

K<sub>a</sub> absorption rate constant, t<sub>1/2</sub>K<sub>a</sub>absorption half-life, K distribution rate constant, t<sub>1/2</sub>K distribution half-life, K'<sub>a</sub> re-absorption rate constant, t<sub>1/2</sub>K<sub>a</sub> re-absorption half-life, K'elimination rate constant, t<sub>1/2</sub>K' elimination half-life, Vd<sub>area</sub> apparent volume of distribution, AUC total area under the plasma ceftriaxone concentration vs. time curve.

**Table 4:** Mean urine concentration (μg ml^−1^) of ceftizoxime (metabolite of ceftriaxone) and ceftriaxone in healthy lactating goats without and with 1 hr pre single dose oral administration of fibrosin (1.9 gm) after single dose intramuscular administration of ceftriaxone at 50 mg kg^−1^

| Time (hr) | Fibrosin treated healthy lactating |  |
| --- | --- | --- |
|  | Ceftriaxone | Ceftizoxime |
| 0 – 24 | $67.91 \pm 9.42$ | BDL |
| 24 – 48 | BDL | BDL |
| 48 – 72 | BDL | BDL |

BDL – Below detectable level

Ceftriaxone at a concentration of 67.91 ± 9.42 μg ml^−1^ was recovered at 24 hr post dosing from urine of the lactating goats. However, neither ceftriaxone nor ceftizoxime (active metabolite of ceftriaxone) could be detected in urine at 48 and 72 hr post dosing.

## Discussion

Single dose oral administration of Fibrosin^®^ (1.9 gm) did not change Icterus index, serum bromosulphophthalein clearance (BSP), serum aspartate transaminase activity (AST) and serum alanine transaminase activity (ALT) significantly (p<0.05) between different days (0, 3 and 7) which indicated that Fibrosin^®^ does not alter normal liver functions.

Plasma concentration of ceftriaxone showed an increasing - decreasing pattern followed by the same increasing- decreasing pattern again in lactating goats following single intramuscular dosing with 1 hr prior oral administration of Fibrosin^®^ (Table 2). Disposition of ceftriaxone exhibited shorter absoption half life (t_½_ka=0.064+0.004 hr) and distribution half life (t_½_k=0.091±0.01 hr) compared to increased reabsorption half life (t_½_ K′a=0.18±0.003 hr) and elimination half life (t½k′=0.43±0.035 hr) with prior oral administration of the polyherbal drug (Table 3). Hepatic clearance of ceftriaxone predominated renal clearance in goat (Sar *et al*, 2008). Ceftriaxone was found to be an inducer of microsomal cytochrome P_450_ in liver and undergoes hydrolysis to produce its active metabolite i.e. ceftizoxime through cleavage of thioether bond present in ceftriaxone (Sar *et al*, 2006). Furthermore, ceftrixone having a molecular weight 554.59 is liable to excrete through bile that may undergo reabsorption in the intestine resulting in its entry into the systemic circulation followed by metabolism in liver (Sar *et al*, 2011; Sar *et al*, 2013) or reabsorption again. Therefore, biphasic increased concentration of ceftriaxone after intramuscular administration in these animals may suggest enterohepatic circulation of ceftriaxone and its intestinal reabsorption in goats.

Ceftriaxone persisted for 3 hr in plasma of lactating goats following single intramuscular dosing at 50 mg kg^−1^with 1 hr prior oral administration of Fibrosin^®^ in the present study while it persisted for 2 hr in lactating goats without the polyherbal drug (Sar *et al*, 2013).The Ka(absorption rate constant) of ceftriaxone in presence of the polyherbal drug was 10.75±0.80 hr^−1^(Table 3) in the present study which increased markedly compared to Ka value of 8.41±0.02 hr^−1^ in healthy goats without oral administration of the polyherbal drug(Sar *et al*, 2013).It indicated that the polyherbal drug has increased the absorption rate of ceftriaxone from intramuscular site as well as the persistence of the drug in the body. Moreover, it can be transpired that Fibrosin^®^, an inhibitor of cytochrome P_450_ oxidase system may cause lesser/negligible metabolism of ceftriaxone through this system leading to longer persistence compared to healthy goats without fibrosin administration.

Ceftriaxone persisted for 2hr in plasma in healthy goats following single intramuscular dosing at 50 mg kg^−1^ and showed a reabsorption half life (t_½_Ka′) of 0.09 ± 0.005 hr (Sar *et al*, 2013). But it persisted for a longer time in fibrosin treated goats which might be due to its enhanced intestinal reabsorption and impaired metabolism in liver compared to healthy goats which is also evidenced by its two fold increased half life of reabsorption (t_½_Ka′=0.18 ± 0.003 hr) (Table 3). Ceftriaxone also showed reabsorption following enterohepatic circulation in healthy goats after intramammary administration of ceftriaxone with 1 hr prior oral dosing of Fibrosin^®^(Sar *et al*, 2011).The drug also showed reabsorption phase in the present study with concomitant administration of the polyherbal drug. Ceftriaxone, a lipid soluble drug having molecular weight of 554.59 is expected to excrete through biliary secretion. Enterohepatic circulation of ceftriaxone may either induce cytochrome P_450_ enzyme of the liver (Sar *et al*, 2006) and subsequently being metabolized or excrete through bile as such. Enhanced reabsorption of ceftriaxone from intestine occurred in healthy lactating goats following intramuscular administration in the present study indicating lesser metabolism of ceftriaxone due to prior oral administration of the polyherbal drug. It was also evidenced by an abruptly increased reabsorption half life with prior oral administration of the polyherbal drug as it was reported to inhibit cytochrome P_450_ enzyme of the liver (Sar *et al*, 2006). Sar *et al*, 2006; could not detect reabsorption of ceftriaxone in goats following single dose intravenous administration of ceftriaxone at 50 mg kg^−1^(a higher dose) without prior oral dosing of the polyherbal drug (Fibrosin^®^) which might be due to optimum induction of cytochrome P_450_ in liver with subsequent rapid metabolism as a result of cent percent bioavailability. One of the functions of liver is secretion of bile which contains pigments and bile salts. Bile salts are formed in the liver from cholesterol and excreted as sodium salts of taurocholic and glycocholic acids after conjugation with taurine and glycine which help in the emulsification of fats in the intestine for absorption of fat soluble vitamins and compounds (Sastry, 1999). Ceftriaxone, being a lipid soluble drug may be absorbed in presence of bile from the intestine. Failure of bile excretion due to hepatic damage was responsible either for accumulation of drug in the liver or interference with the further absorption of ceftriaxone resulting in absence of reabsorption phase in the disposition of the drug in hepatopathic goats (Sar *et al*, 2013).Ceftriaxone showed major hepatic clearence in goats which was evidenced by its higher C1_H_ value (Sar *et al*, 2008) and intestinal reabsorption of ceftriaxone occurred in the present study following intramuscular administration which indicated that the polyherbal drug did not interfere with bile secretion from liver and caused enhanced excretion of ceftriaxone through bile. The t_½_k ′(elimination half life) of ceftriaxone was 0.43 ± 0.035 hr with prior oral administration of the polyherbal drug in the present study. But it decreased drastically compared to t_½_k′ value of 0.76 ± 0.09 hr in healthy goats following intramuscular dosing of ceftriaxone at the same dose rate without Fibrosin^®^ administration (Sar *et al*, 2013) which may be due to enhanced excretion of the drug through bile. No reabsorption phase could be observed in disposition of ceftriaxone in previous studies by other research workers in healthy animals after intramuscular administration which might be due to collection of blood samples at comparatively longer intervals. Blood samples from fibrosin treated lactating goats were collected at comparatively shorter intervals from our previous experience during the study (Sar *et al*, 2013).

Ceftizoxime could not be detected in urine of fibrosin treated healthy goats which further suggested negligible metabolism of ceftriaxone leading to non-availability of its metabolite, ceftizoxime. It is reported that ceftriaxone induces cytochrome P_450_ in liver and undergoes hydrolysis to form the major active metabolite i.e. ceftizoxime following single dose intravenous and intramuscular administration at 50 mg Kg^−1^ in goats which was excreted at higher concentration in urine up to 72 hr of collection (Sar *et al*, 2006; Sar *et al*, 2013). Oral Fibrosin^®^ administration may decrease the content of cytochrome P_450_ which is responsible for impaired metabolism of ceftriaxone in healthy goats resulting in non availability of its metabolite in urine. Ceftriaxone is mixed function oxidase system inducer causing more formation of its own metabolite (ceftizoxime) which is expected to excrete through urine in healthy goats. But ceftriaxone has lesser/no effect on MFO system in presence of fibrosin resulting in negligible amount of metabolite formation which might be the reason of non availability of ceftizoxime in urine of fibrosin treated goats. Ceftriaxone is inducer of hepatic microsomal enzyme systems while fibrosinis the inhibitor of the enzyme and combined therapy induced enzyme activity (Sar *et al*, 2006). Interestingly, combined therapy of intramuscular ceftriaxone injection and oral Fibrosin^®^(polyherbal drug) in the present study showed negligible induction of MFO system resulting in non availability of its active metabolite i.e. ceftizoxime in urine. This might be due to lesser bioavailability of ceftriaxone following intramuscular administration for which the inhibition of cytochrome P_450_ suppressed induction of cytochrome P_450_ by ceftriaxone. Either ceftizoxime or ceftriaxone also could not be detected in milk following intramuscular administration of ceftriaxone at 50 mg kg ^−1^ with 1 hr prior oral dosing of the polyherbal drug (Sar *el al*, 2014).

Intramuscular administration of ceftriaxone with concomitant oral administration of Fibrosin^®^ may be recommended particularly as persistence of the antibiotic is increased in plasma with zero level milk residue and lesser degree of elimination for shorter duration in urine that may subside public health hazard and lesser chances of development of antimicrobial resistance to both ceftriaxone and ceftizoxime.

The reabsorption of ceftriaxone following intramuscular administration even with concurrent administration of cytochrome P_450_ inhibitor should be taken into account for calculating therapeutic dosage regimen, drug interactions and drug monitoring in different disease conditions. Moreover, Fibrosin^®^ (the polyherbal drug) could be used as a representative cytochrome P_450_ inhibitor for pharmacokinetic interaction study of ceftriaxone particularly as it did not interfere with bile secretion and liver function tests like icterus index, serum bromosulfophthalein clearance (BSP), serum aspartate transaminase activity (AST), serum alanine transaminase activity (ALT) but inhibited cytochrome P_450_oxidase system in liver.

## Materials and Methods

### Drugs and Chemicals

Ceftriaxone (analytical grade, purity ≥ 90 %;Estral Pharmaceutical Industries, Vadadora, Gujarat, India) was used as test drug. Ceftizoxime (analytical grade, purity ≥ 90 %) was obtained from GlaxoSmithkline Pharmaceuticals Ltd, Nashik, India. Other chemicals were obtained from Sigma Aldrich, USA.Fibrosin^®^ (Legend Remedies Pvt. Ltd.,Vadodara, India) is a polyherbaldrug used for concomitant therapy with antibiotics in clinical and sub-clinical mastitis. It is composed of Kanchanar-gugal (gummy substances and resins of *Bauhinia variegata*Linn.), Chitrak-mula(root of *Plumbagozeylanica*). Punar-navastaka(flower of *Triaanthemamonogyna*), + Trifala(fruit of *TerminaliabelericaRetz* + fruit of *Terminaliachebula*Retz.+fruit of *Phylanthusamblica*) and Apamarga(whole plant of *Achyranthesaspera*Linn.). The actual composition of Fibrosin^®^ is trade secret of Legend Remedies Pvt. Ltd.,Vadadora, India. The active principles present in these plants were presented in our earlier studies (Sar *et al*, 2006; Sar *et al*, 2011). Fibrosin^®^ was used as ctochome P_450_ inhibitor for pharmacokinetic interaction study with intramuscular ceftriaxone injection.

### Animals

A total of 6 apparently healthy lactating goats weighing between 10-12 kg of approximately 1½-2 years of age were used in this experiment. The animals were caged individually in custom made metabolic cages (stainless steel) during experimental period. Artificial lighting facility was provided with 12 hour dark period. Animals were stall-fed and the standard feed as well as water was provided *ad libitum*.The composition of the feed was 2 parts wheat husk, 1 part groundnut cake, 1 part crushed gram, 1 part crushed maize, and 2 parts greens. The animals were dewormed with a single oral dose of rafoxanide at 7.5 mg kg^−1^ body weight 30 days prior to the onset of study. Before the start of the experiment, the animals were acclimatized for 7 days.

### Experimental design

The effects of Fibrosin^®^ in liver of six healthy lactating goats were assessed by performing icterus index, serum bromosulphophthalein clearance (BSP), serum aspartate transaminase activity (AST), serum alanine transaminase activity (ALT). For this study, a half bolus of Fibrosin^®^ (1.9 gm) was administered orally making suspension in 50 ml distilled water to each lactating goat and the tests were performed collecting serum before oral dosing of Fibrosin^®^ and on day 3 and 7 after Fibrosin^®^ administration.

For pharmacokinetic interaction study, a single dose of ceftriaxone dissolving in 5 ml of distilled water was administered at 50 mg kg^−1^ body weight to each animal deep at thigh muscle. A half bolus of fibrosin (1.9 gm) was administered orally to each goat before 1 hr of ceftriaxone administration making suspension in 50 ml distilled water. The same animals were utilized for pharmacokinetic interaction study after allowing a washing period of 1month.

### Collection of samples

Blood samples werecollected from jugular vein at ‘0’ and at 0.08, 0.16, 0.25, 0.33, 0.50, 0.66, 0.83, 1, 2, 3 and 4 hr post dosing of intramuscular ceftriaxone injection. Plasma was then separated and utilized for estimation of drug concentration.

Urine samples were collected at 0, 24, 48 and 72 hr post dosing for analysis of ceftriaxone/ceftizoxime.

### Biochemical tests

#### Bromosulphophthalein (BSP) clearance test

The test was carried out in the serum of goats before administration of fibrosin^®^(1.9gm) and on day 3 and 7 and bromosulphophthalein (BSP) clearance (Varley *et al*, 1975) was expressed as t_½_min (elimination half life).

#### Icterus index

The test was carried out in the serum of goats^17^at ‘0’ (pre dosing of fibrosin^®^), day 3 and 7 following oral administration of fibrosin^®^ and was expressed as unit.

#### Estimation of serum AST and ALT activity

Serum AST (aspartate aminotransferase) and ALT (alanine aminotransferase) activities were measured according to the method of Yatazidis (Yatazidis, 1960).

#### Analysis of ceftriaxone and ceftizoxime

Extraction of ceftriaxone as well as its metabolite, ceftizoxime from plasma and urine was done which were subsequently analyzed by HPLC. Plasma or urine (0.5 ml) was taken in a test tube, 0.5 ml of mobile phase was added to it. After that 2 ml of the mixture containing 70% methanol, sodium acetate (30% of 0.1M) added to the test tube to make a total volume of 3 ml and pH (5.2) was adjusted by using HPLC grade acetic acid (1-2 drops). The mixture was vortexed for 30 minute and incubated at 4^0^C for 15 min. The whole aliquot was centrifuged at 3000 rpm for 15 min. and supernatant was collected after passing through a filter paper (Whatman No. 1 Rankem).The supernatant (20 μl) was analysed by SHIMADZU LC-20AT liquid chromatogram coupled with Photo Diode-Array detector (UV-VIS) attached with computer SPD MXA 10 software. A 5μ Luna C18 (2); 250 x 4.6 mm (RP) column was used with the flow rate of 1.0 ml/min and 254 nm wavelength. The mobile phase was prepared according to the method of USP (United States Pharmacopoeia) for analysis of ceftriaxone. Standard and sample (20μl) were injected into the injector port of liquid chromatography with the first and last being the standard. The drugs were estimated after comparing with external standard. Calibration was done every time by a standard stock solution of 20 ppm of the test drugs of analytical grade prepared in distilled water. Recovery of ceftriaxone/ceftizoxime from plasma and urine was carried out in vitro by adding known quantities of ceftriaxone/ceftizoxime to give final concentrations of 0.005, 0.01, 0.05, 0.1, 0.25, 0.50,1,2,5,10 and 20μg ml^−1^.The limit of detection for both the drugs was 0.01 μg ml^−1^. The retention times of ceftriaxone and ceftizoxime were 9.549 and 4.504, respectively which showed inter-day variations.

#### Pharmacokinetic analysis

The concentration of ceftriaxone in plasma of lactating healthy goats at different time intervals was plotted on a semilogarithmic paper and the pharmacokinetic parameters were analysed by non-compartmental method. The initial increased concentration was considered as absorption phase (K_a_) which was followed by decreased concentration and considered as distribution phase (K). The concentration increased again considered as reabsorption phase (K_a_) followed by decreased concentration considered as elimination phase (K). All these half-lives were calculated from the same graph and AUC (area under curve) was determined based on trapezoid method. The other pharmacokinetic parameters were determined as per the method of Baggot (Baggot, 1977). The pharmacokinetic parameters such as K, K′, K_a_, K′_a_ are indicated for K distribution rate constant, K′ elimination rate constant, K_a_ absorption rate constant, K′_a_ reabsorption rate constant in healthy lactating goats in which pharmacokinetic profile of ceftriaxone showed a distinct reabsorption phase.

#### Statistical analysis

The results were expressed as Mean ± Standard error (S.E.). The data were analyzed statistically using general linear model with univariate data in SPSS 10.0 version of software (students’s t test).

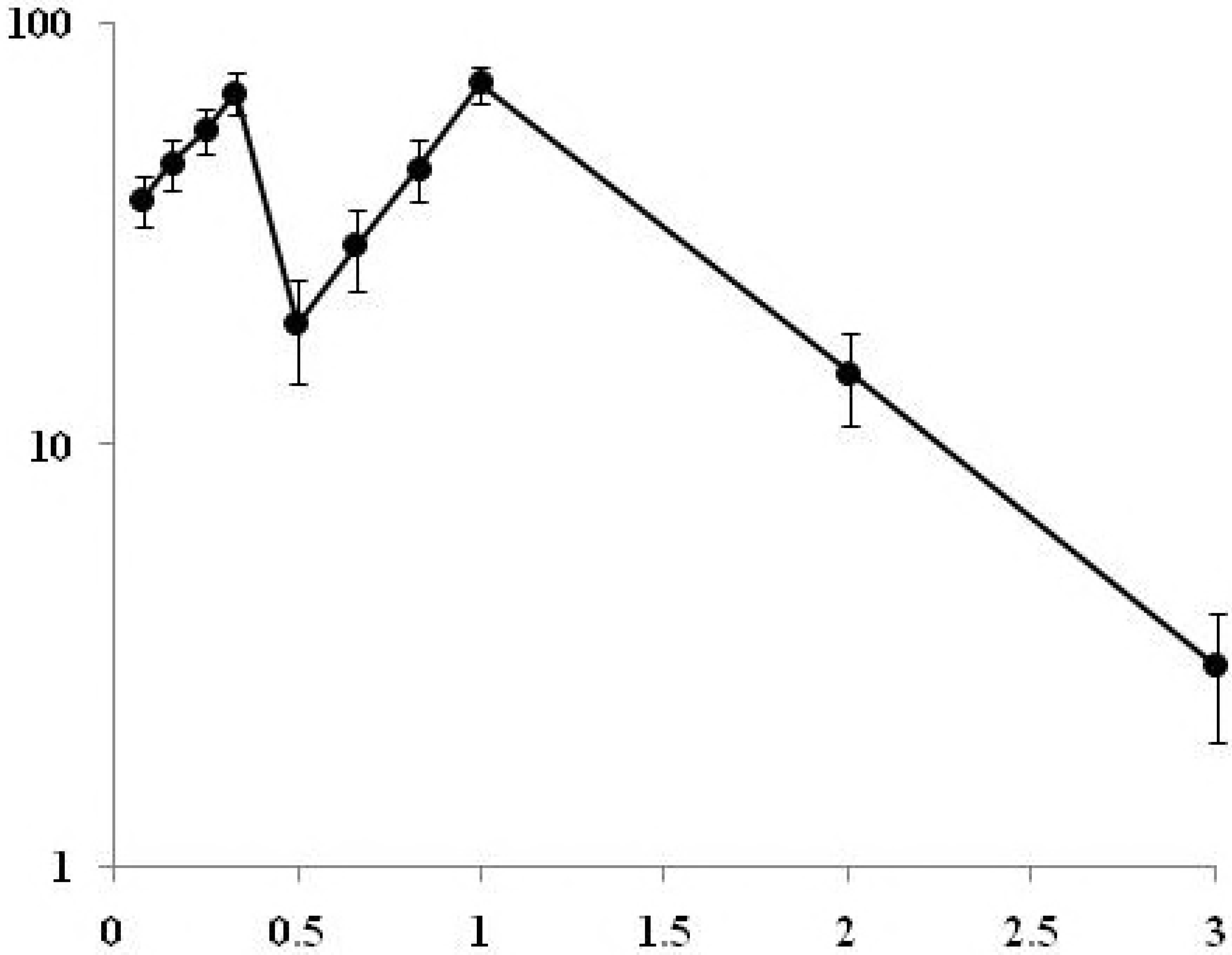

## Acknowledgements

The authors provide sincere gratitude to the honourable Vice Chancellor, West Bengal University of Animal and Fishery Sciences for necessary facilities and infrastructure. The authors are thankful to Dr. Amlan Kumar Patra, Department of Animal Nutrition, WBUAFS for statistical analysis and editing the manuscript.

### Author Contributions

TKS conducted the experiment, IS and RNH wrote and edited the manuscript, RB helped to modify estimation method, TKM designed the experiment

### Conflict of interest

The authors declare that they have no conflict of interest.

### Compliance with Ethical Standards

No external fund was received by any of the author for the work. The authors would like to disclose no potential conflicts of interests. All applicable institutional guidelines for the care and use of animals were followed.

